# Maternal tamoxifen treatment expands the macrophage population of early mouse embryos

**DOI:** 10.1101/296749

**Authors:** Rocío Rojo, Kristin A. Sauter, Lucas Lefevre, David A. Hume, Clare Pridans

**Affiliations:** The Roslin Institute, University of Edinburgh, Easter Bush EH25 9RG, UK.; Current address: UK Dementia Research Institute, University of Edinburgh, Edinburgh EH16 4SB, UK.; Current address: Mater Research Institute-University of Queensland, Translational Research Institute, Woolloongabba QLD 4104, Australia.; Current address: Medical Research Council Centre for Inflammation Research at the University of Edinburgh, Edinburgh EH16 4TJ, UK

## Abstract

Several different transgenic tamoxifen-inducible cre reporter lines have been used to analyse the contribution of embryonic precursors to the development of the mononuclear phagocyte system in mice. Here we show that tamoxifen treatment of the mother at 8.5dpc with doses commonly-used in lineage trace studies produces a 4-5-fold expansion of the embryonic leukocyte populations by 10.5dpc, detected in whole mounts of embryos using a *Csf1r* reporter gene or separately by expression of *Csf1r, Itgam* (CD11b), *Adgre1* (F4/80) or *Ptprc* (CD45) mRNA. These findings indicate that tamoxifen cannot be considered a neutral agonist in macrophage lineage trace studies.

**Summary sentence:** Treatment of pregnant mice with tamoxifen in early gestation produces a large expansion of the embryonic macrophage population.

## Introduction

The mononuclear phagocyte system (MPS) is a family of cells that includes committed progenitors, blood monocytes and tissue macrophages. Mononuclear phagocytes are found in every adult tissue (1, 2) where they adapt to local environments to perform specific functions (3-7). The original definition of the MPS proposed that tissue macrophages were continuously replaced by blood monocytes (8). The emerging alternative view is that most tissue macrophages, with exceptions such as dermis and gut, are seeded during embryonic development from yolk sac progenitors and maintained mainly by self-renewal (6, 9-12). This view depends in part upon the interpretation of fate-mapping studies in which tamoxifen treatment of the mother induces recombination and activation of a reporter gene in the embryo. The key assumption in lineage-trace studies is that a single treatment with tamoxifen produces recombination only in cells that are present in the embryo within the narrow time window prior to the onset of definitive hematopoiesis (9). However, Epelman *et al* (10), suggested that tamoxifen might persist in tissues and initiate recombination in cells at later stages as they infiltrate those tissues. Where this was tested directly in adult mice in an islet transplantation model, there was clear evidence of such persistence (11). Ye *et al.* (12) highlighted persistent accumulation of tamoxifen and nuclear translocation of Cre-ER(T2) at least 2 months beyond a single pulse, as well as direct effects of tamoxifen on adipogenesis. Similarly, a single low-dose tamoxifen treatment of juvenile males produced long term adverse effect on multiple endocrine systems (13).

A second assumption in the use of tamoxifen-inducible reporters is that the drug does not alter the differentiation of monocyte-macrophage cells in the embryo, based in part upon the reported lack of expression of the estrogen receptors in foetal haematopoietic progenitors (14). However, this assumption is questioned by evidence of the direct effects of estrogen on macrophage gene expression and self-renewal *in vivo* and *in vitro* (15). Patel *et al* (13) noted that >40% of published studies that use tamoxifen to generate conditional mutations lack controls for the impact of tamoxifen. We therefore decided to investigate whether tamoxifen treatment of the mother, at doses used in reported lineage trace studies, alters the development of embryonic macrophage populations.

## Materials and Methods

Approval was obtained from The Roslin Institute and The University of Edinburgh Protocols and Ethics Committees under the authority of a UK Home Office Project License under the regulations of the Animals (Scientific Procedures) Act 1986. Adult (>8 weeks) wild type CD1 females and *Csf1r*-ECFP^+^ (MacBlue) males were used for this study. Each female was placed overnight with one male in a cage. Detection of a vaginal plug the following morning was considered as 0.5 days postcoitum (dpc). At 8.5 dpc, mice were orally gavaged with vehicle (2.5% ethanol in sunflower oil (Sigma S5007)) or 5 mg of 4-Hydroxytamoxifen (Sigma-Aldrich, H6278), resuspended in vehicle. A naïve group was untreated.

At 10.5 dpc females were culled by cervical dislocation and embryos were dissected. Embryos were imaged using a StereoLumar v12 (Zeiss) dissection microscope. Excitation and emission wavelengths used to detect ECFP signal were 426-446 nm and 460-500nm, respectively.

For mRNA quantitation, total RNA from 10.5 dpc mouse embryos was isolated using RNeasy Mini Plus kit (Qiagen). The quantity and quality of total RNA were controlled by using the NanoDrop ND-1000 spectrophotometer (Thermo Fisher Scientific) and the 2200 TapeStation (Agilent) respectively. The RNA Integrity Number (RIN) values were all >7. cDNA was synthesised from 1 ug total RNA with oligo(dT)20 and random primers (3:1, v/v) using Superscript III reverse transcriptase (Thermo Fisher Scientific) according to the manufacturer’s instructions. For quantitative PCR, cDNA was amplified with Fast SYBR Green Master Mix using the 7500 fast Real Time PCR system (Applied Biosystems, Thermo Fisher Scientific). The oligonucleotides spanning an intron were designed using Primer3 web version 4.0.0.

The oligonucleotides used were:

*Hprt* fwd 5’- GCGATGATGAACCAGGTTATGA-3’

*Hprt* rev 5’- CCTTCATGACATCTCGAGCAAG-3’

*Csf1* fwd 5’- AGTATTGCCAAGGAGGTGTCAG-3’

*Csf1* rev 5’- ATCTGGCATGAAGTCTCCATTT-3’ (16).

*Csf1r* fwd 5’- CAGTTCAGAGTGATGTGTGGTC-3’

*Csf1r* rev 5’- CTTGTTGTTCACTAGGATGCCG-3’

*Adgre* (Emr1) fwd 5’- TCTGGGGAGCTTACGATGGA-3’

*Adgre* (Emr1) rev 5’- ACAGCAGGAAGGTGGCTATG-3’.

*Ptprc* (CD45) fwd 5’- AACAGAGCCTCAGCCTACACTC-3’

*Ptprc* (CD45) rev 5’- TAGACACCAGGTCTTTGGGTTC-3’

*Itgam* (CD11b) fwd 5’- CTGTCACACTGAGCAGAAATCC-3’

*Itgam* (CD11b) rev 5’- AGGATGGAAGGTCACACTGAAT-3’

*Kit* fwd 5’- GAAAGTGATGTCTGGTCCTATGG-3’

*Kit* rev 5’- CCTTGATCATCTTGTAGAACTTGG-3’.

Primer efficiency was validated with a standard curve of four serial dilution points (efficiency ranging between 95% and 105%). Data were normalised using the dCq model (17).

## Results and discussion

The MacBlue transgenic mice have a co-insertion of a modified *Csf1r* promoter driving the artificial transcription factor, Gal4-VP16, alongside a Gal4-dependent UAS-ECFP reporter. In many adult organs, the ECFP reporter provides a marker for recently-arrived monocytes, which are ECFP-positive, where most adult tissue macrophages are ECFP-negative (18, 19). In the embryo, the ECFP marker is detectable in the earliest macrophage-like cells in Reichert’s membrane and retained on the majority of embryonic macrophages throughout development (19). Because the ECFP marker is so bright, it can be detected readily in whole mounts of the embryo and provides a unique global overview of macrophage development.

We injected 4-OH tamoxifen into four pregnant wild-type female mice mated to MacBlue positive males at 8.5dpc using standard protocols, doses and vehicle described by others (e.g. (9, 20-22)). At 10.5dpc, we removed the embryos and compared the pattern of ECFP expression. Two controls were untreated, and two were treated with the vehicle. An increase in ECFP-positive cells in response to treatment was evident throughout the body, notably concentrated in the ventricles of the developing brain, where cells expressing *Csf1r* mRNA are concentrated at 10.5dpc (23). There was also a smaller, but detectable, effect of the injection of the vehicle alone (Figure 1).

**Figure 1.**
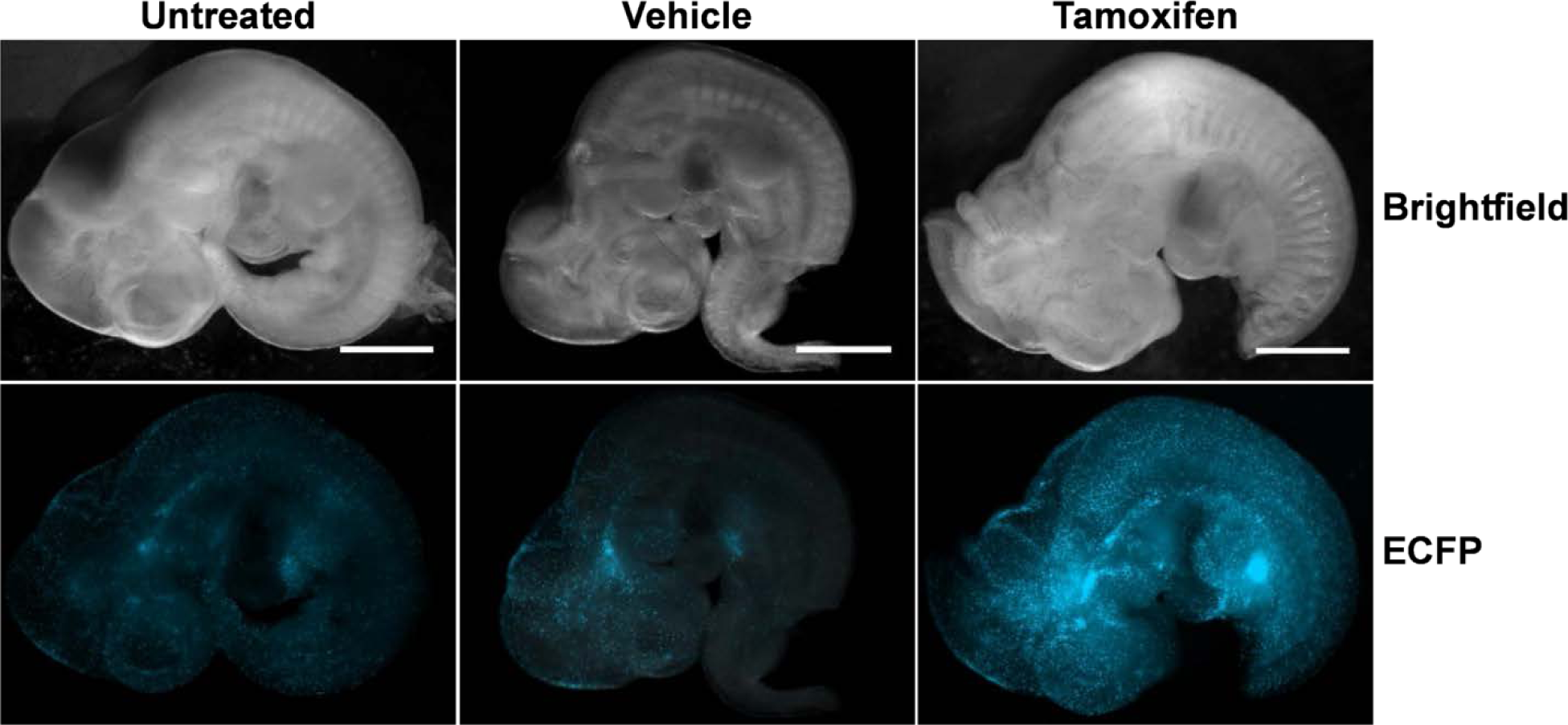
Visualisation of the impact of maternal tamoxifen treatment at 8.5dpc on embryonic macrophages. Pregnant wild-type female mice mated to MacBlue^+^ males were injected with vehicle or 4-OH tamoxifen at 8.5dpc. Untreated dams were not injected. Embryos were collected at 10.5dpc and examined for ECFP expression. Images are representative of 3 (untreated), 4 (vehicle) and 8 (tamoxifen) embryos from two separate litters. Scale bar=500µm.

To assess the difference, we used ImageJ to quantify embryo size and fluorescence intensity (Figure 2). ECFP fluorescence was increased around 4-fold compared to untreated or control vehicle-treated embryos. Since macrophages are a major source of somatic growth factors, including Insulin-like growth factor 1 (IGF1) in embryonic and postnatal development (24), the increased size of the embryo could be a consequence of increased macrophage numbers.

**Figure 2.**
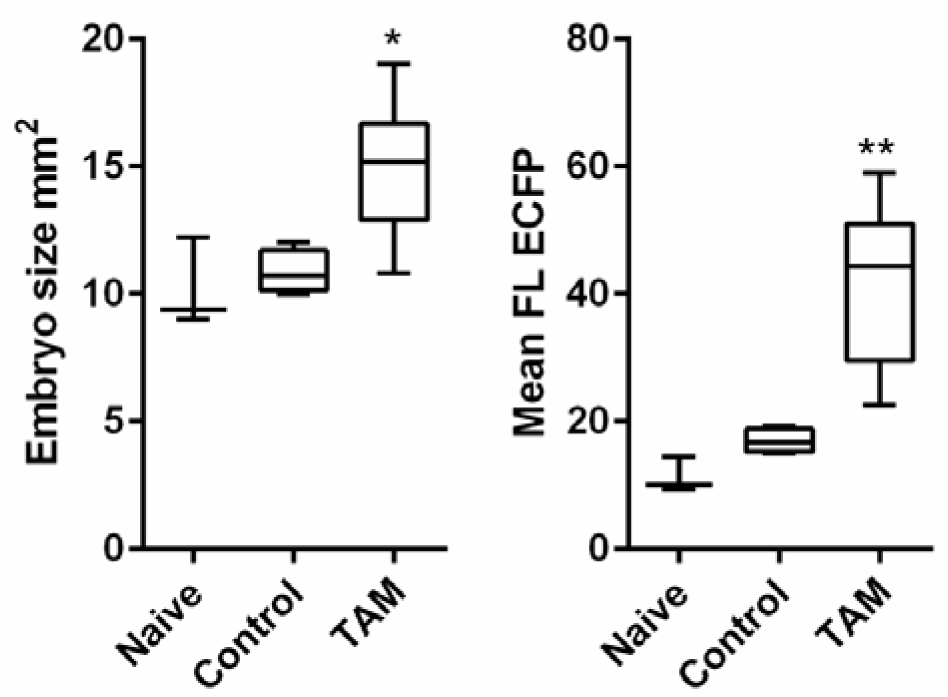
Quantitation of the impact of maternal tamoxifen treatment at 8.5dpc on embryonic macrophages. Images from embryos of untreated (naïve) mothers, or treated with vehicle (control) or 4-OH tamoxifen (TAM), were quantified using Image J as described in Materials and Methods. n = 3 naïve and 4 control embryos, and 8 embryos from TAM-treated dams from two separate litters. * *P* = 0.0213 TAM relative to naïve (size) and ** *P* = 0.0028 TAM relative to naïve (FL; fluorescence) via a student’s t-test.

The impact of maternal tamoxifen treatment on embryonic macrophage development was examined further in a separate cohort of pregnant mice by analysis of the expression of markers associated with leukocytes or macrophages (Figure 3). Like the *Csf1r* reporter gene, *Csf1r* mRNA expression is restricted to macrophages in the developing embryo (23). Accordingly, the *Csf1r* promoter has been used to drive macrophage-specific inducible recombinase expression in many lineage trace studies (21, 22, 25). A cluster of 120 macrophage-associated transcripts, including *Adgre1, Itgam, Cd68*, *Sirpa*, *Fcgr1* (Cd64) and *Mertk*, was strongly correlated with *Csf1r* in a time course of mouse embryonic development (26). The coordinated expression of genes within this cluster provided a surrogate for the expansion of tissue macrophage numbers in multiple tissues in the developing embryo. The expression of *Csf1r*, and of two other macrophage-associated transcripts, *Adgre1* (F4/80) and *Itgam* (Cd11b) mRNA, was increased 4- to 5-fold in the embryos from tamoxifen-treated mothers. The pan-leukocyte marker, *Ptprc* (CD45) was also elevated to the same extent as the macrophage markers. At this developmental stage, the large majority of leukocytes in the whole embryo are macrophages (23) and *Ptprc* expression parallels *Csf1r* (26).

**Figure 3.**
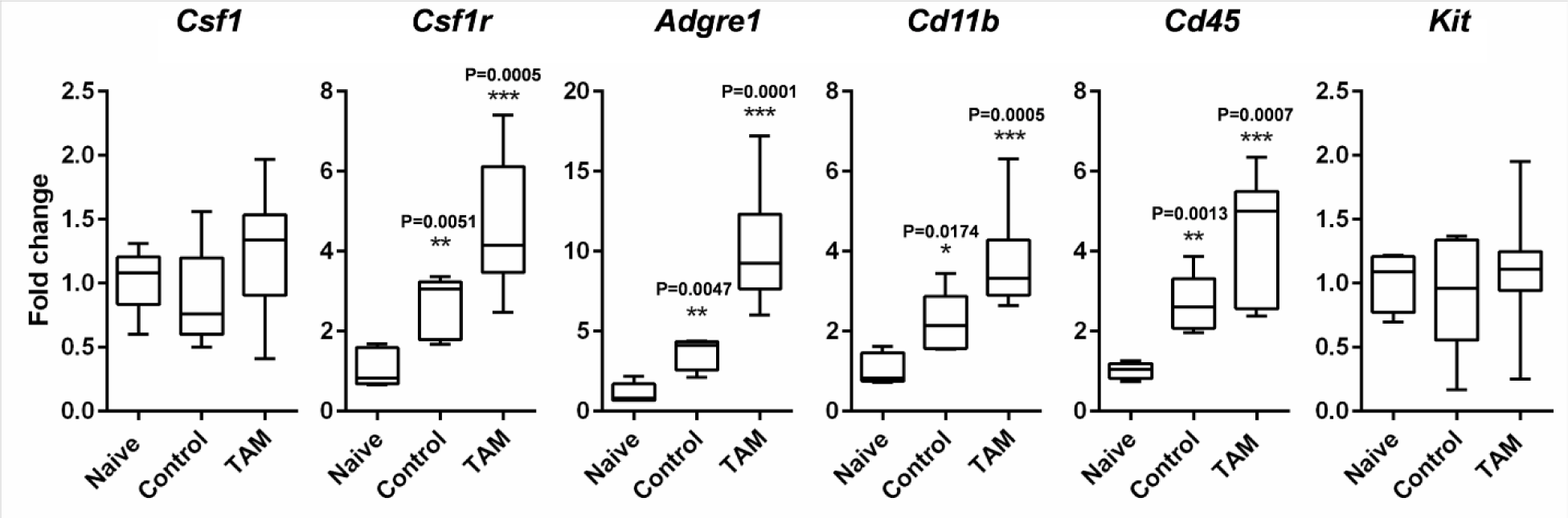
The impact of maternal tamoxifen treatment at 8.5dpc on gene expression in the embryo at 10.5dpc. mRNA was isolated from embryos from untreated mothers (naïve), or treated with vehicle (Control) or 4-OH tamoxifen (TAM), and mRNA quantified as described in Materials and Methods. Results for naïve and control are from 5 embryos. Results for the TAM-treated animals are from 9 embryos from two separate litters. P values (relative to naïve) are indicated on graphs and were calculated via a student’s t-test.

In each case, there was a smaller impact of treatment with vehicle compared to naïve mice. This is a second control that is not commonly performed. The vehicle is 2.5% ethanol in sunflower oil. It is possible that either of the components of the vehicle is pharmacologically active. Perhaps more likely, the oral gavage *per se* may produce a stress response that can also impact the embryo. Walker *et al* (27) demonstrated a significant activation of the hypothalamic-pituitary-adrenal axis in response to oral gavage in mice. Whilst we are not aware of direct evidence for impacts on early macrophage development, there is a large literature on the impact of maternal stress on the immune system of offspring (28).

In view of the known sensitivity of *Csf1* mRNA to hormonal stimulation (29) and the impact of maternal anti-CSF1R treatment to reduce embryonic macrophage numbers (21) we considered the possibility that the treatment(s) might induce *Csf1* mRNA in the embryo. However, although *Csf1* mRNA was readily detected, it did not vary amongst treatment groups (Figure 3). Finally, we also examined expression of *Kit*, which is expressed by erythro-myeloid progenitor cells in the yolk sac (21). Results obtained using a tamoxifen-inducible lineage trace based upon *Kit* suggested that most adult tissue macrophages are derived from definitive haematopoietic stem cells (30). By contrast to the macrophage-lineage specific genes, *Kit* mRNA was not increased in response to tamoxifen, suggesting that the treatment does not act directly to expand the progenitor pool.

Depending upon the nature of the estrogen receptor complex in target cells, tamoxifen can be either an agonist or an antagonist of estrogen receptor signaling (31). Samokhvalov et al. (9) based their original lineage trace protocol on the reported lack of estrogen receptors in foetal hematopoietic progenitors. However, tamoxifen might equally act on the mother, for example regulating growth factor availability across the placenta. The lineage trace study indicating that myb-independent cells generated in the yolk sac contribute to the development of adult macrophages (22) included co-administration of progesterone, to prevent complications of tamoxifen administration late in pregnancy as proposed by earlier authors (32). Estrogen and progesterone together have been shown to produce substantial increases in expression of *Csf1* mRNA in the uterus. *Csf1* in turn regulates development of the placenta (29). The production of *Csf2* (GM-CSF) mRNA by uterine epithelial cells is also highly-regulated by ovarian steroid hormones (33). Alternatively, like most cancer chemotherapeutic agents, tamoxifen has multiple side effects mediated by interactions with targets other than estrogen receptors. For example, tamoxifen is a potent inhibitor of the lysosomal enzyme, acid ceramidase, which in turn regulates the levels of pro-apoptotic ceramide and mitogenic sphingosine-1-phosphate (34).

Actively phagocytic macrophages derived from the yolk sac proliferate extensively as they infiltrate tissues. We speculate that the expansion of yolk sac-derived macrophage populations in response to tamoxifen would produce an over-estimate of their contribution to adult macrophage populations. If tamoxifen administered at 8.5dpc is retained in tissues (10), it could also compromise the onset of definitive haematopoiesis in the liver at 10.5dpc. Haematopoietic stem cells are estrogen-responsive and estrogen receptor signalling is required for the expansion of stem cell self-renewal and erythropoiesis that occurs in female mice during pregnancy (35). Treatment of adult mice with tamoxifen at doses used herein produced a marked reduction in bone marrow cellularity, and depletion of haematopoietic stem cells (36).

There are fate-mapping studies using promoters such as *Runx1, S100a4, Cx3cr1, Kit* and *Flt3* that do not involve the use of tamoxifen. Rather than supporting an exclusive foetal origin of tissue macrophages, many such studies indicate progressive contributions from the recruited monocyte progeny of definitive hematopoietic stem cells (Reviewed by (10)). For example, a recent detailed study of peritoneal macrophages indicated their progressive replacement by monocyte-derived cells with age (37). In the developing chick, where *in ovo* cellular transplantation is possible, macrophages derived from transplanted yolk sac progenitors are replaced by the time of hatch whereas those derived from transplanted bone marrow are retained in adults (38).

A compromise view is that macrophages in tissues occupy specific anatomical niches or territories which can be filled either by monocyte infiltration or by local proliferation of resident macrophages depending upon circumstance, opportunity and host genotype (1, 2, 39). Based upon the impacts of treatment with CSF1, or with anti-CSF1R, the local availability of this key growth factor, regulated by macrophages, may be one determinant of niche occupancy (2, 39). This compromise is compatible with the original descriptions of the mononuclear phagocyte system, almost 40 years ago, which clearly recognised the dual origin of tissue macrophages (40-42).

